# Intraflagellar transport during the assembly of flagella of different length in *Trypanosoma brucei* isolated from tsetse flies

**DOI:** 10.1101/2020.05.14.095216

**Authors:** Eloïse Bertiaux, Adeline Mallet, Brice Rotureau, Philippe Bastin

## Abstract

Multicellular organisms assemble cilia and flagella of precise lengths differing from one cell to another, yet little is known about the mechanisms governing these differences. Similarly, protists assemble flagella of different lengths according to the stage of their life cycle. This is the case of *Trypanosoma brucei* that assembles flagella of 3 to 30 µm during its development in the tsetse fly. It provides an opportunity to examine how cells naturally modulate organelle length. Flagella are constructed by addition of new blocks at their distal end via intraflagellar transport (IFT). Immunofluorescence assays, 3-D electron microscopy and live cell imaging revealed that IFT was present in all life cycle stages. IFT proteins are concentrated at the base, IFT trains are located along doublets 3-4 & 7-8 and travel bidirectionally in the flagellum. Quantitative analysis demonstrated that the total amount of IFT proteins correlates with the length of the flagellum. Surprisingly, the shortest flagellum exhibited a supplementary large amount of dynamic IFT material at its distal end. The contribution of IFT and other factors to the regulation of flagellum length is discussed.

**Summary statement:** This work investigated the assembly of flagella of different length during the development of *Trypanosoma brucei* in the tsetse fly, revealing a direct correlation between the amount of intraflagellar transport proteins and flagellum length.

## Introduction

Cilia and flagella are typical eukaryotic organelles found in numerous organisms. They share a similar architecture based on a cylinder of 9 doublet microtubules (with a central pair for most motile organelles) but their length varies exhaustively from one organism to the other and even from one cell type to another within the same organism. Several proteins involved in ciliary length control have been identified but the actual mechanisms are still debated (Broekhuis et al., 2013; Goehring and Hyman, 2012; Ishikawa and Marshall, 2017; Keeling et al., 2016; Lechtreck et al., 2017). These organelles are constructed by addition of new subunits at their distal end (Bastin et al., 1999; Johnson and Rosenbaum, 1992). Since flagella do not contain ribosomes, a transport process called intraflagellar transport (IFT) is required to bring precursors from the cytoplasm of the cell body to the assembly site (Craft et al., 2015; Kozminski et al., 1993; Wren et al., 2013). IFT is the movement of large protein arrays (or trains) made of 20 IFT proteins driven by molecular motors: kinesin II for anterograde transport and cytoplasmic dynein 2 for retrograde transport (Prevo et al., 2017). It is conserved in all organisms assembling cilia and flagella on a protrusion of the cell body (Avidor-Reiss and Leroux, 2015).

Because of its central function in flagellum construction, IFT is a prime candidate for flagellum length regulation. In *Chlamydomonas reinhardtii* and *Caenorhabditis elegans*, cilia and flagella undergo turnover of tubulin at their distal end and their length has been proposed to be the result of a balance between assembly and disassembly rates (Hao et al., 2011; Marshall and Rosenbaum, 2001). These rates can be modulated by changes in train length (Engel et al., 2009) and frequency (Ludington et al., 2013) and/or by loading of flagellar precursors on trains (Craft et al., 2015; Wren et al., 2013). The molecular mechanisms controlling these parameters remain to be elucidated, with protein phosphorylation emerging as a potential candidate (Liang et al., 2018; Wang et al., 2019). The majority of these studies have been performed in the green algae *Chlamydomonas*, which is a powerful model for the study of flagellum assembly. It constructs two flagella of equivalent length that can be manipulated by various genetic and biochemical approaches (Dutcher, 2014).

Although these manipulations offer great tools to investigate the mechanisms involved in flagellum length control, they might not necessarily reflect the main means by which cells control the length of their organelle in natural conditions. In that context, protists are interesting organisms as they can naturally assemble flagella of different lengths according to their stage of development. Some of them even possess several types of flagella in the same cell, leading to sophisticated system to discriminate both in terms of construction and function (Bertiaux and Bastin, 2020; Bertiaux et al., 2018b; McInally et al., 2019). *Trypanosoma brucei* is a typical example and exhibits multiple life cycle stages characterised by flagella ranging from 3 µm to more than 30 µm (Rotureau et al., 2011; Sharma et al., 2008; Van Den Abbeele et al., 1999)(Fig. 1). It therefore offers the possibility to investigate which parameters are modulated to build flagella of different length.

**Fig. 1.**
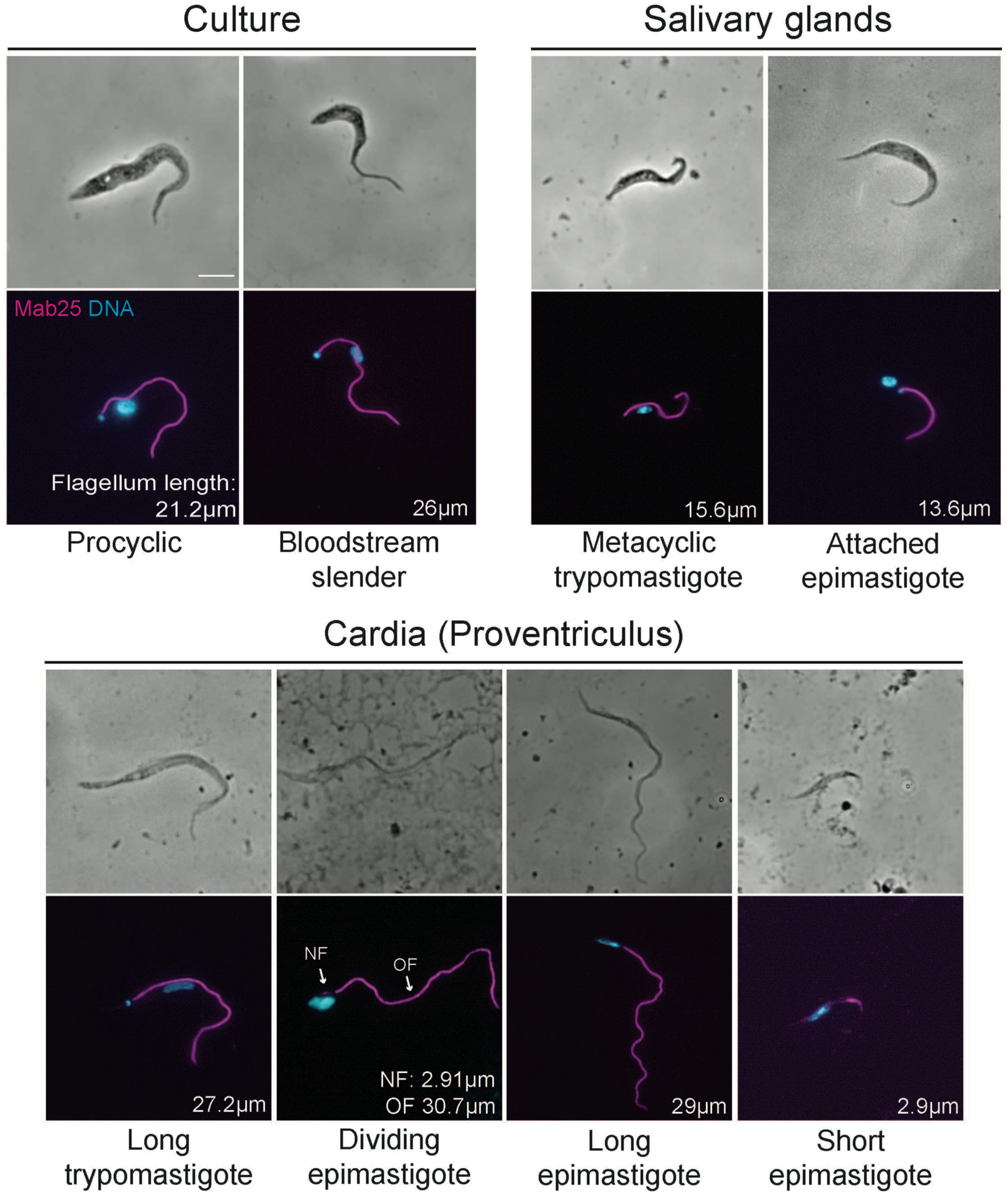
Evolution of flagellum length during the parasite cycle. The main morphological stages of the *T. brucei* parasite cycle encountered in the tsetse fly vector are presented, as well as the two forms that can be grown in culture. Parasites were fixed with cold methanol and stained with DAPI (cyan) and the Mab25 antibody (magenta) that recognises an axoneme component (Dacheux et al., 2012). Flagellum length was measured using the Mab25 marker and is indicated for each stage. Data is from (Rotureau et al., 2011), with the exception of dividing epimastigote cells that were specifically analysed in this study (n=37). The scale bar represents 5 µm.

Trypanosomes are parasites infecting mammals via the bite of the tsetse fly. In the insect, they first develop in the midgut as the procyclic stage. They possess a single flagellum that is attached to the cell body and that emerges from the posterior end of the cell. This configuration is termed trypomastigote (Fig. 1). This stage is proliferative and can be grown in culture. Trypanosomes then transform into the so-called mesocyclic stage (Rotureau et al., 2011; Sharma et al., 2008; Van Den Abbeele et al., 1999), which is defined by the presence of a longer flagellum (Fig. 1). However, since there are no molecular markers and that these cells seem to be able to revert at the procyclic stage in certain conditions (Van Den Abbeele et al., 1999), we prefer to use the term long trypomastigote (Fig. 1). These parasites migrate to the cardia (proventriculus) and concomitantly undergo exhaustive morphogenetic modifications to become epimastigote (Lemos et al., 2020; Sharma et al., 2008). The epimastigote configuration is defined by the emergence of the flagellum at the anterior part of the cell (Fig. 1). These cells divide asymmetrically, producing a long epimastigote daughter that inherits the old flagellum (∼30 µm) and a short epimastigote daughter with a tiny flagellum (∼3 µm)(Fig. 1). Once parasites have reached the salivary glands, they attach to the epithelium via large outgrowths of the flagellar membrane (Tetley and Vickerman, 1985) and then switch to the trypomastigote configuration during the emergence of the metacyclic stage (Fig. 1) that is released in the saliva and ready for transmission. The mechanisms governing the formation of these different types of flagella are unknown. Investigation has been complicated by the absence of cultivation system for most of these stages and therefore requires analysis of parasites extracted from infected tsetse flies.

All *IFT* genes are conserved in trypanosomes (Morga and Bastin, 2013) and functional investigation by RNAi knockdown revealed that they are essential for flagellum construction (Absalon et al., 2008; Davidge et al., 2006; Franklin and Ullu, 2010; Kohl et al., 2003). IFT trafficking itself has been investigated exhaustively in the procyclic stage that is amenable to cell culture and manipulation. The frequency and speed of anterograde (∼1 train/s and ∼2 µm/s) and retrograde trafficking (∼2-3 trains/s and ∼5 µm/s) have been determined using live analysis of the movement of various IFT proteins such as IFT22 (Adhiambo et al., 2009), IFT25 (Huet et al., 2019), IFT27 (Huet et al., 2014), IFT52 (Buisson et al., 2013) and IFT81 (Bhogaraju et al., 2013) as well as several subunits of the dynein motors (Blisnick et al., 2014).

Here, we investigated IFT in several natural life cycle stages of *T. brucei* that exhibit flagella of different lengths. We mostly focused on trypanosomes present in the anterior midgut and the cardia of infected tsetse flies where the most extreme differences in flagellum length have been reported (Rotureau et al., 2011; Sharma et al., 2008; Van Den Abbeele et al., 1999). Conventional transmission electron microscopy and focused ion beam scanning electron microscopy (FIB-SEM) revealed the presence of IFT trains in the flagellum of trypanosomes found in the cardia. Immunofluorescence assays using two different antibodies and monitoring of live trypanosomes expressing a fluorescent IFT protein showed the presence of IFT in all stages analysed, from procyclic in the midgut to metacyclic in the saliva. IFT trafficking was clearly visible in all cells observed, however video recording turned out to be extremely challenging due to the high flagellum beating rate and the difficulty to immobilise cells without damage. Nevertheless, quantification of IFT amount could be achieved in fixed cells for all stages and live IFT tracking was successful in certain stages. In both cases, the total amount of IFT material was directly correlated to flagellum length. By contrast, the IFT concentration per unit of flagellum length appeared constant. The only exception was the tiny flagellum of short epimastigote that contains a supplementary and dynamic pool of IFT proteins at its distal end that could contribute to its elongation in the next developmental stage. Finally, several experiments were performed in an effort to understand why flagellum length is so short at that particular stage.

## Materials and Methods

### Trypanosome cell lines and cultures

Derivatives of *T. brucei brucei* strain AnTat 1.1E Paris procyclic strain (Le Ray et al., 1977) were cultured at 27°C in SDM79 medium supplemented with hemin, 8mM of glycerol and 10% foetal calf serum (Brun and Schonenberger, 1979). For bloodstream form trypanosomes, strain 427 was cultured in complete HMI-11 medium at 37°C in 5% CO2 (Kooy, Hirumi et al. 1989). IFT imaging in live cells was carried out with a cell line expressing a TdTomato::IFT81 fusion produced from its endogenous locus in AnTat1.1E PCF cells. The first 500 nucleotides of the *IFT81* gene (Gene DB number Tb927.10.2640) were chemically synthesised (GeneCust, Luxembourg) and cloned in frame with the TdTomato gene within the HindIII and ApaI sites of the p2675 vector (Bertiaux et al., 2018b; Kelly et al., 2007). The construct was linearized within the *IFT81* sequence with the enzyme XcmI and nucleofected in the AnTat1.1E PCF cell line, leading to integration by homologous recombination in the *IFT81* endogenous locus and to the expression of the full-length coding sequence of IFT81 fused to TdTomato.

### Tsetse fly maintenance, infection and dissection

Tsetse flies (*Glossina morsitans morsitans*) were maintained, infected and dissected at the Institut Pasteur as previously described (Rotureau et al., 2011). Teneral males were collected 24 to 48 h post-eclosion and fed through a silicone membrane with 6-9×10^6^ parasites/ml in SDM-79 medium supplemented with 10% FCS, 8mM glycerol and 10mM glutathione for their first meal (MacLeod et al., 2007). Flies were infected with the *T. brucei brucei* AnTat 1.1E Paris wild type (Le Ray et al., 1977) and AnTat 1.1E IFT81::TdT strains. Flies were then maintained in Roubaud cages for one month at 27°C and 50% humidity and fed three times a week with mechanically defibrinated sheep blood. Flies were starved for at least 48h before being individually dissected 28 days after ingestion of the infected meal. In our colony conditions, the average midgut infection rates usually obtained with AnTat 1.1E Paris wild-type strain are 50-60% in the midgut and 10-15% in the salivary glands. Salivary glands were first rapidly dissected into a drop of PBS or culture medium. The whole tsetse alimentary tract was then dissected and arranged lengthways for assessing the parasite presence. The cardia and anterior midgut were physically separated from the posterior midgut in a distinct PBS drop. Tissues were dilacerated to allow parasites to spread in PBS; parasites were recovered and treated for further experiments no more than 15 minutes after dissection.

### Electron microscopy

For electron microscopy sample preparation, the entire digestive tracts of 11 flies were placed in a drop of 0.1 M cacodylate buffer (pH 7.2) and fixed in 2.5% glutaraldehyde (Sigma-Aldrich), 4% paraformaldehyde. Entire cardia were then separated from the digestive tract and transferred to 1.5 ml Eppendorf tubes containing 500 µl of 2.5% glutaraldehyde, 4% paraformaldehyde in 0.1 M cacodylate buffer (pH 7.2) for 1 or 2 h at 4° C. Fixed samples were washed three times by the addition of fresh 0.1 M cacodylate buffer (pH 7.2) buffer and post-fixed in 1% osmium (EMS) in 0.1 M cacodylate buffer (pH 7.2) enriched with 1.5% potassium ferrocyanide (Sigma-Aldrich) for 50 min in the dark under agitation. Samples were gradually dehydrated in acetone (Sigma-Aldrich) series from 50% to 100%. Cardia were oriented along longitudinal or transversal axes and embedded in PolyBed812 resin (EMS) hard protocol (Luft, 1961), followed by polymerization for 48 h at 60° C. After embedding, 500 nm semi-thin sections were performed and stained with toluidine blue to find the region of interest containing trypanosomes. Semi-thin sections were used like a map to mill the sample precisely in the region of interest by FIB. For TEM analysis, ultra-thin sections (70 nm) were obtained using an ultramicrotome (Leica EM UC7). Sections were post-stained 40 minutes with 4% uranyl acetate and 10 minutes with 3% lead citrate and observed using a FEI Tecnai T12 120kV. For FIB-SEM, the embedded sample was coated with 20nm of Au/Pd and directly placed in the chamber of a FESEM Zeiss Auriga microscope equipped with a CrossBeam workstation (Zeiss) and acquired using ATLAS 3D software (Zeiss) as described (Bertiaux et al., 2018a).

### Immunofluorescence

Cultured parasites were washed twice in PBS and spread directly into poly-L-lysine coated slides. The slides with cultured parasites or parasites isolated from tsetse flies were air-dried for 10 min, fixed in methanol at -20°C for 30 s and rehydrated for 10 min in PBS. For immuno-detection, slides were incubated with primary antibodies diluted in PBS with 0.1% Bovine Serum Albumin (BSA) for 1 h at 37°C in humid chamber. The antibodies used were the Mab25 mouse monoclonal antibody recognizing TbSAXO1, a protein found all along the trypanosome axoneme (Dacheux et al., 2012; Pradel et al., 2006), an anti-IFT172 mouse monoclonal antibody diluted at 1/200 (Absalon et al., 2008), an anti-IFT22 mouse polyclonal antiserum diluted at 1/200 (Adhiambo et al., 2009), an anti-FLAM8 rabbit polyclonal antibody diluted at 1/500 (kind gift of Paul McKean, Lancaster University). Three washes of 10 min were performed and the secondary antibody diluted in PBS with 0.1% BSA was added on the slides. Subclass-specific secondary antibodies coupled to Alexa 488 and Cy3 (1/400; Jackson ImmunoResearch Laboratories, West Grove, PA) were used for double labelling. After an incubation of 45 min, slides were washed three times in PBS for 10 min and DAPI (4′,6-diamidino-2-phenylindole) (2 µg/µl) was added to stain DNA. Slides were mounted with coverslips using ProLong antifade reagent (Invitrogen). Sample observation was performed using a DMI4000 microscope equipped with a 100X NA 1.4 objective (Leica, Wetzlar, Germany) and images were captured with an ORCA-03G Hamamatsu camera. Pictures were analysed using the ImageJ 1.47g13 software (National Institutes of Health, Bethesda, MD). For presentation purposes and only after analysis, images were merged using Photoshop CS6 (Adobe).

### Live imaging of parasites isolated from tsetse fly

Fly dissections were performed in SDM-79 medium. The midgut, cardia and salivary glands were rapidly separated in different drops. After a visual inspection to confirm the presence of parasite, each organ was kept in 15µL of SDM-79 medium supplemented with serum in Eppendorf tubes. To increase parasite density 6 organs were pooled in each tube and dilacerated. A SDM-79 solution containing 4% Agar was heat-liquefied and progressively cooled for being mixed with the medium containing the freshly isolated parasites (1:1). For live video microscopy, the solution was spread on a slide, covered with a coverslip and rapidly observed at room temperature under the DMI4000 microscope. Videos were acquired using an Evolve 512 EMCCD Camera (Photometrics, Tucson, AZ) driven by the Metaview acquisition software (Molecular Probes, Sunnyvale, CA). IFT trafficking was recorded at 100 ms per frame during 30 seconds. Kymographs were extracted and analysed using Quia software as described previously (Buisson et al., 2013; Chenouard et al., 2010).

### FRAP analysis

For FRAP analysis of cells expressing Tdt::IFT81, an Axiovert 200 inverted microscope (Zeiss) equipped with an oil immersion objective (magnification x100 with 1.4 numerical aperture) and a spinning disk confocal head (CSU22, Yokogawa) was used (Buisson et al., 2013). Movies were acquired using Volocity software with an EMCCD camera (C-9100, Hamamatsu) operating in streaming mode. The samples were kept at 27°C using a thermo-controlled chamber. Sequences of 30 s were acquired at an exposure time of 0.1 s per frame.

### Inhibition of cell division

Flies were dissected in SDM-79 medium and after visual inspection; three positive proventriculi were pooled and resuspended in 500 µL of SDM-79 supplemented with hemin, 10% fetal bovine serum and 10mM glycerol in 24-wells plates. For inhibiting cell division, teniposide (Sigma SML0609), a topoisomerase II inhibitor was dissolved in DMSO and added to isolated parasites from tsetse flies at a final concentration of 10 mM for 24 hours (Bertiaux et al., 2018b; Robinson and Gull, 1991). In the control wells, the same volume of DMSO alone was added (6.3µL). Parasites were spread on poly-lysine slides and allowed to sediment for 30 minutes at 27°C before being fixed and processed for immunofluorescence.

## Results

Different methods are available for probing IFT: visualisation by electron microscopy, detection of IFT proteins by immunofluorescence in fixed cells and recording of IFT trafficking in live cells following expression of IFT proteins fused to fluorescent proteins. All these approaches were attempted here.

### IFT trains are detected in trypanosomes developing in the tsetse fly

IFT trains were first identified by correlative light and electron microscopy (Kozminski et al., 1995) as electron-dense particles sandwiched between microtubule doublets and the flagellar membrane of *Chlamydomonas* (Ringo, 1967). In *T. brucei*, similar densities are visible in sections of bloodstream form trypanosomes isolated from rats (Vickerman, 1962) or of procyclic form parasites grown in culture (Sherwin and Gull, 1989).

To look for IFT trains in various stages of *T. brucei*, we have infected tsetse flies with the AnTat 1.1E strain and dissected the cardia, where cells with the shortest and longest flagella are encountered (Fig. 1). The cardia is a relatively large organ (Fig. S1A) and although parasites can be found at fairly high densities, it is essential to probe the sample for presence of trypanosomes before performing sections for electron microscopy. First, we did semi-thin sections (500 nm) and stained them with toluidine blue (Fig. S1B). Once regions with sufficient parasite density were detected, ultrathin sections were carried out. Observation at low magnification by transmission electron microscopy revealed the presence of multiple parasites in various orientations (Fig. 2A,B). Classic trypanosome organelles were frequently observed such as the nucleus, the mitochondrion or acidocalcisomes, as well as the characteristic subpellicular microtubules (Fig. 2A,B). Sections through the base of the flagellum revealed the flagellar pocket and its collar, from which the flagellum emerges at the cell surface (Fig. 2A,B). Multiple flagellum sections were visible, with the presence of the canonical axoneme and of the associated paraflagellar rod (PFR). These images confirm that trypanosomes present in the cardia display the typical cytological organisation as reported in other stages. Examples of flagellum cross sections at higher magnification are presented at Figures 2C-E. The 9 doublets can be numbered unambiguously thanks to the fixed orientation of the central pair and the associated PFR (Fig. 2C). Densities equivalent to IFT trains were frequently observed between the membrane and the microtubules (Fig. 2C-E) and they were mostly encountered in proximity of doublets 3-4 (Fig. 2E) or 7-8 (Fig. 2C,D), as observed in procyclic cells (Absalon et al., 2008; Bertiaux et al., 2018a).

**Fig. 2.**
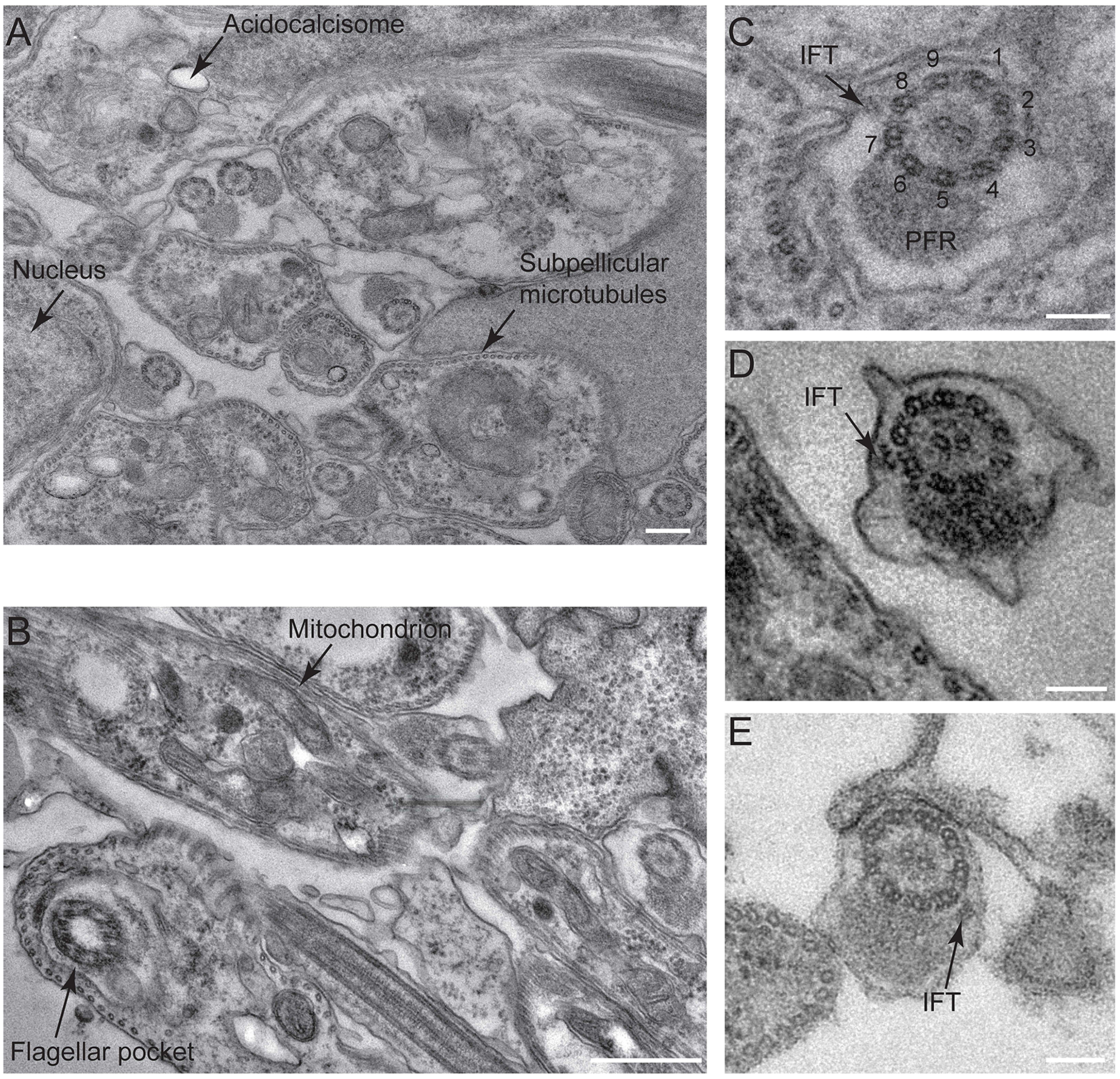
IFT trains are present in the flagellum of parasites found in the cardia. (A,B) Sections at low magnification of fields showing tsetse fly cardia infected by *T. brucei* parasites. Typical organelles such as the nucleus, an acidocalcisome, the subpellicular microtubules (A), the flagellar pocket or the mitochondrion (B) are indicated. Multiple sections through flagella are visible. (C-E) Higher magnification of flagellum sections with IFT particles positioned close to doublets 7-8 (C and D) or doublet 4 (E). The numbering of doublets and the paraflagellar rod (PFR) are indicated on image C. The scale bars represents 200 nm (A), 500 nm (B) and 100 nm (C-E).

Thin sectioning does not allow the identification of the parasite stage and we therefore turned to FIB-SEM for further analysis. Using the same approach as above with semi-thin sectioning, we identified a region that was suitable for milling by FIB (Fig. S2C). Several series were acquired with a 10 nm Z-resolution. In the volume presented, a total of 722 sections were performed (Video S1 and Fig. 3). On that sample, the epithelium of the cardia is present on the right-hand side and trypanosomes can be found towards the bottom left. Three images corresponding to slices 225, 298 and 374 are presented (Fig. 3A). The flagellum highlighted at Fig. 3 is visible on 470 consecutive slices (Video S1). The presence of the flagellum attachment zone (FAZ, Fig. 3B) allows the identification of the cell body it belongs to. Although the base of the flagellum is not in the volume, the nucleus is visible on the first ∼200 slices with the attached flagellum, indicating that this cell is not at the epimastigote stage. Based on the diameter of the cell body, this cell is likely to transit between the long trypommastigote and the epimastigote stage (Lemos et al., 2020; Sharma et al., 2008). Looking at that flagellum more closely, a train is visible in proximity of doublet 8 at image 114 and stretches until section 315 (Video S1, Fig. 3B, left and central panels). A second train is present on that same flagellum but this time in vicinity of doublet 4 on image 359 until image 396 (Video S1, Fig. 3B, right panel). This distribution is similar to that reported in procyclic trypanosomes grown in culture (Bertiaux et al., 2018a). Despite careful analysis, we could not find a short epimastigote in the two volumes analysed (Lemos et al., 2020). Although FIB-SEM offers excellent conditions to identify IFT trains and determine their positioning, finding all the stages of interest is like looking for a needle in a haystack and we therefore turned towards light microscopy for quantitative analysis of IFT using immunofluorescence with anti-IFT antibodies and live cell analysis upon expression of IFT fluorescent fusion proteins.

**Fig. 3.**
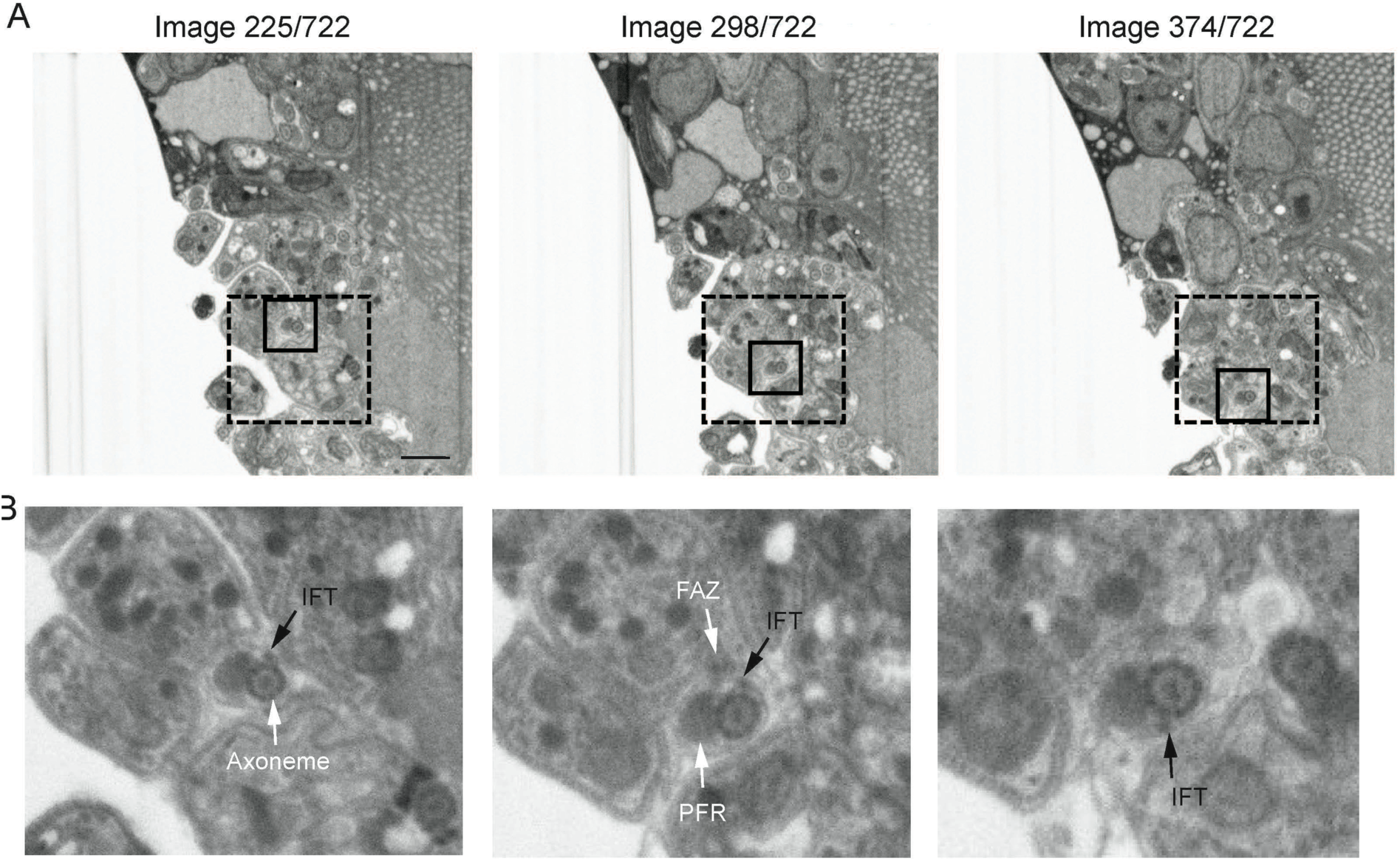
IFT train distribution by 3-D electron microscopy. (A) A portion of the cardia infected with trypanosomes was analysed by FIB-SEM. Three images extracted from Video S1 are presented at low magnification images (position in the stack is indicated on top of each image). Several trypanosomes are visible on the left-hand side while tsetse tissues are on the right-hand side. The dotted squares indicate the region of interest that is shown in Video S1. The full square indicated the portion that is showed at higher magnification in panel B. (B) Each image comes from the field shown above. IFT particles are indicated, with a long train found on doublet 8 (left and centre images) and a shorter one present on doublet 4 (right image). The presence of the FAZ confirms which cell body the flagellum belongs to. The full sequence is presented at Video S1.

### IFT amounts correlate with flagellum length at all stages of the parasite cycle

In procyclic trypanosomes in culture, IFT proteins are progressively recruited during flagellum elongation with a direct correlation between IFT total amounts and actual flagellum length, whilst the frequency of injection and the concentration of IFT proteins per unit of length remain unchanged (Bertiaux et al., 2018b). Similar features have been reported recently in the different pairs of flagella in *Giardia* (McInally et al., 2019). To evaluate the possible link between IFT amounts and flagellum length when trypanosomes assemble different flagella during their life cycle, the distribution of IFT proteins was investigated in fixed cells coming either from infected flies or from cultured bloodstream and procyclic stages. Immunofluorescence assays (IFA) with antibodies raised against IFT172 (Fig. 4A)(Absalon et al., 2008) or IFT22/RABL5 (Fig. S2)(Adhiambo et al., 2009) were performed in combination with the axonemal marker Mab25 (Pradel et al., 2006). For all parasite stages, IFT proteins were present as a succession of diffuse spots all along the length of the flagellum, with a brighter signal at the base present in almost all stages except in short epimastigote cells (Fig. 4A and Fig. S2).

**Fig. 4:**
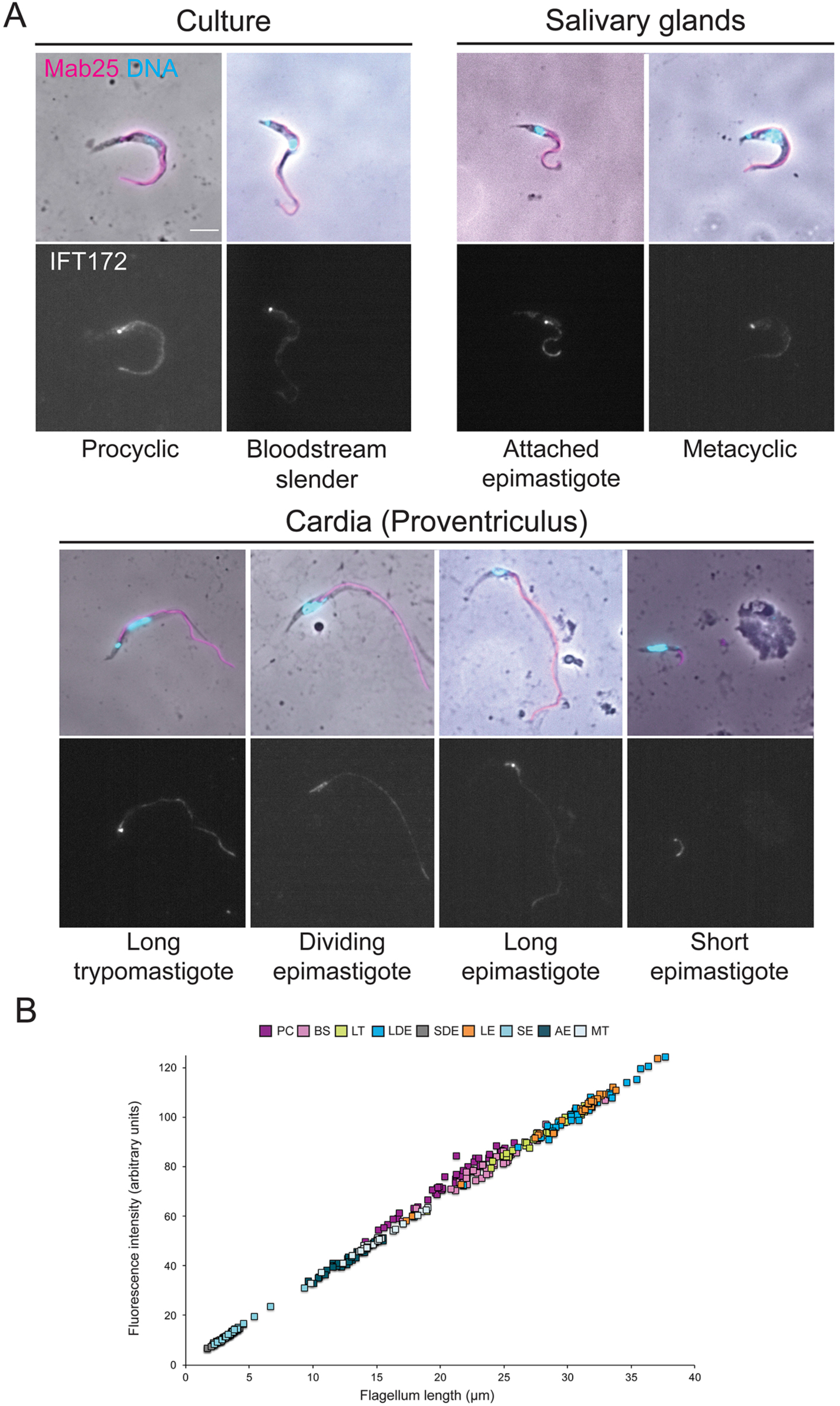
IFT172 distribution during the *T. brucei* parasite cycle. (A) Parasites isolated from tsetse flies or grown in culture were fixed in cold methanol and stained with the Mab25 antibody to detect the axoneme (magenta) and the anti-IFT172 antibody (white). The top panels show the phase-contrast images merged with DAPI (cyan) and Mab25 signals (magenta). The bottom ones show the IFT172 fluorescent signal (white). Scale bar: 5µm. (B) Quantification of the total amount of IFT172 fluorescent signal in the flagellum normalized to flagellum length for each stage of the parasite cycle. The ROI was defined by the axonemal marker and used to measure the flagellum length. The fluorescence intensity directly reflects the total amount of IFT172 proteins present in the flagellum compartment. n=35 cells per stage. BSF: Slender bloodstream form, PC: Procyclic form, LT: Long trypomastigote, LDE: Long dividing epimastigote, SDE: Short dividing epimastigote, LE: Long epimastigote, SE: Short epimastigote, AE: Attached epimastigote, MT: Metacyclic trypomastigote.

To quantify the total amount of IFT22 or IFT172 protein per flagellum, a region of interest was defined using the Mab25 axonemal marker (shown in magenta in the figures). The total amount of fluorescence, corresponding to the total amount of IFT proteins present in the flagellum, was plotted against flagellum length. For both IFT proteins, a direct correlation between the total amount of IFT proteins and the length of the corresponding flagellum was found (Fig. 4B and Fig. S1B). These data suggest that the total amount of IFT protein correlates with the length of the flagellum at all stages of the parasite cycle.

### IFT trafficking displays similar rates and frequency during all the parasite cycle

The methanol fixation classically used for IFA could lead to the loss of some IFT material and therefore biased the results. Moreover, IFA only provides static information while IFT is a dynamic process. To circumvent these potential issues, the distribution of IFT proteins was examined in live trypanosomes expressing a fusion protein between the IFT-B protein IFT81 and the red fluorescent Tandem Tomato protein (TdT)(Bertiaux et al., 2018b). This fusion protein is expressed upon endogenous tagging under the control of the 3’ untranslated region (UTR) of the *IFT81* gene (Bhogaraju et al., 2013). The cell line expressing this fusion protein was fed to tsetse flies that were dissected one month later. The TdT::IFT81 protein was detected in live trypanosomes at all the different stages of the parasite cycle: procyclic cells in the midgut; parasites at the long trypomastigote, dividing epimastigote, long and short epimastigote stages in the cardia and attached epimastigote form and metacyclic form parasites in the salivary glands (Fig. 5A & Videos S2-S4). In all cases, a higher concentration of IFT proteins was detected at the base of the flagellum and IFT trafficking was visible without ambiguity (Fig. 5A & Videos S2-S4). The only major difference in IFT protein distribution was noticed in short epimastigote cells (see below).

**Fig. 5.**
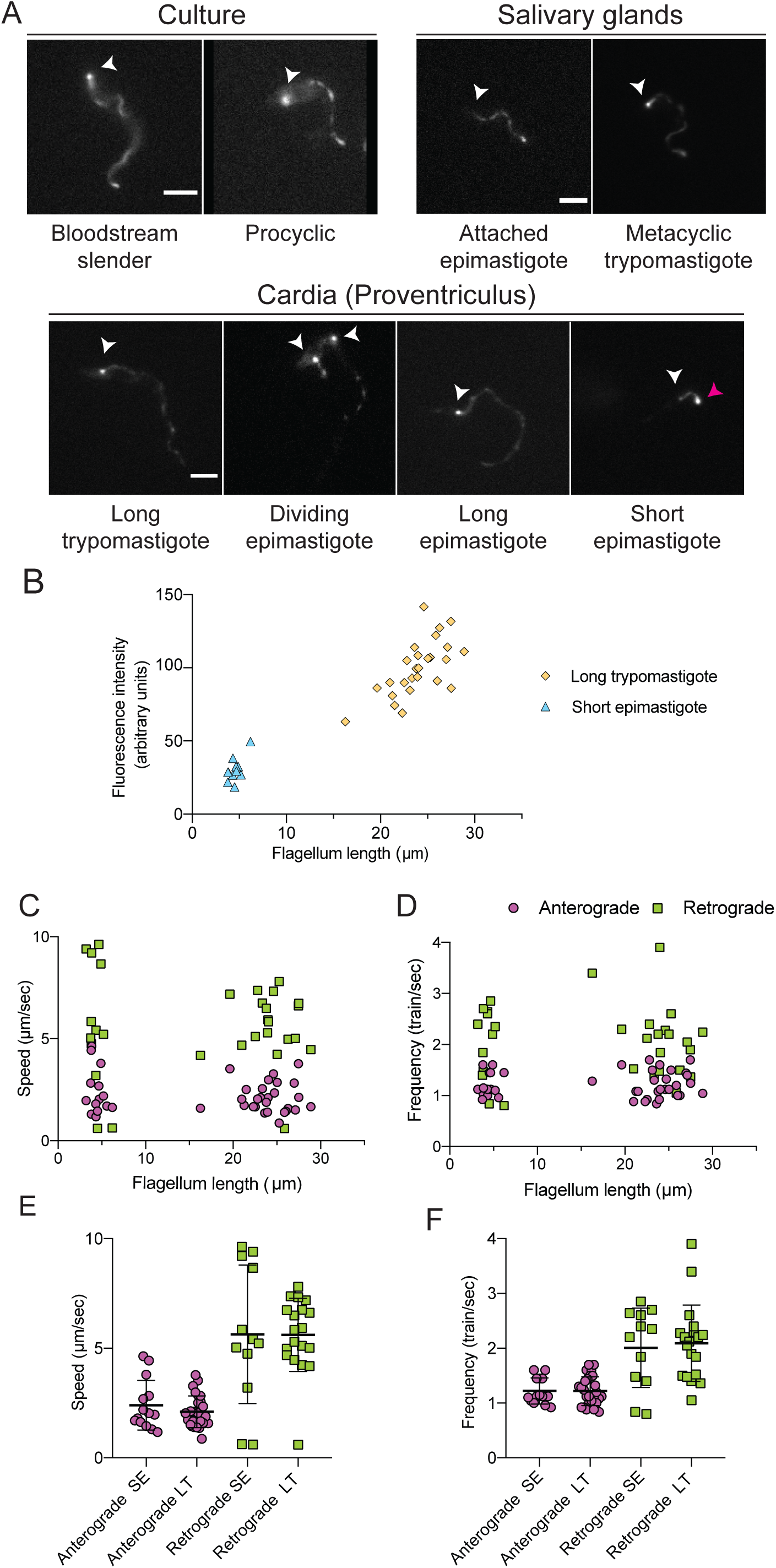
Variation of IFT81 trafficking during the *T. brucei* parasite cycle. (A) Still images extracted from movies of trypanosomes expressing a TdTomato::IFT81 from the endogenous locus at the indicated stages of the parasite cycle either from culture forms or extracted from infected tsetse flies. White arrowheads indicate the base of the flagellum and the magenta arrowhead points at the flagellum tip of a short epimastigote cell. Corresponding videos are presented as Video S2 (culture stages), Video S3 (parasites from the salivary glands) and Video S4 (parasites from the cardia). (B) The total TdT::IFT81 fluorescence intensity in flagella of short epimastigote (blue triangles) and long trypomastigote cells (yellow diamonds) was calculated and plotted according to the length of the corresponding flagella. (C) IFT speeds (anterograde transport, magenta circles; retrograde transport, green squares) in the flagella of short epimastigote and long trypomastigote cells were calculated and plotted according to the respective length of the flagella. Retrograde transport is more difficult to detect and data were therefore only incorporated when the signal was sufficiently intense and reliable. (D) IFT particle frequencies (anterograde transport, magenta circles; retrograde transport, green squares) in the flagellum of short epimastigote cells and long trypomastigote cells were calculated and plotted according to the respective length of the flagella. (E) IFT particle speeds (anterograde transport, magenta circles; retrograde transport, green squares) in the flagella of short epimastigote (SE) and long trypomastigote (LT) cells were determined using kymograph analysis. (F) IFT particle frequencies (anterograde transport, magenta circles; retrograde transport, green squares) in the flagellum of short epimastigote (SE) cells and long trypomastigote (LT) cells were calculated using kymograph analysis.

IFT quantification requires visualisation of the full-length flagellum in the same plane of focus. To restrict flagellum beating, trypanosomes isolated from the cardia of infected flies were mixed with low melting point agar. This could immobilise most procyclic and some long trypomastigote or some short epimastigote cells. Unfortunately, long epimastigote or dividing epimastigote cells remained too motile in these conditions, and all other attempts to immobilise them were unsuccessful, precluding a quantitative analysis for these two stages. Live imaging was therefore only feasible for long trypomastigote (flagellum length between 20 to 30 µm) and short epimastigote cells (flagellum length ∼5 µm). The total amounts of TdT::IFT81 in the flagellum were quantified in live cells of these two stages by using the first image of each movie, and the results were plotted according to flagellum length (Fig. 5B). This demonstrated the direct correlation between these two parameters in live cells, in agreement with the IFA data (Fig. 4B & Fig. S1).

Although the total amount of IFT proteins appeared proportional to the length of the flagellum, modifications of IFT speed and frequency cannot be excluded and could have an impact on the flagellum growth rate (Bertiaux et al., 2018b). Therefore, TdT::IFT81 trafficking was investigated in details with kymograph analyses (Buisson et al., 2013) in cells with a short (short epimastigote) or a long flagellum (procyclic and long trypomastigote forms). In the flagellum of short epimastigote cells, anterograde trains travelled at 2.4 ± 1.3 µm.s^-1^ and retrograde trains at 5.6 ± 3.1 µm.s^-1^ (Fig. 5C & Fig. 5E, n=13). In the flagellum of long trypomastigote or procyclic cells, anterograde trains travelled at 2.1± 0.7 µm.s^-1^ and retrograde trains at 5.6 ± 1.7 µm.s^-1^ (Fig. 5C & Fig. 5E, n=26). Next, the IFT frequency was evaluated: in the short epimastigote flagellum, the frequency of anterograde IFT was 1.2 ± 0.4 trains per second and 2.0 ± 0.7 trains per second for retrograde IFT (Fig. 5D & Fig. 5F). In the long trypomastigote / procyclic flagellum, the anterograde IFT frequency was similar with 1.22 ± 0.3 trains/s while the retrograde IFT frequency was 2.1 ± 0.1 trains/s (Fig. 5D & Fig. 5F). There were no statistical differences of speed and frequency between the two stages based on a t-Student test. It should be noted that these speed and frequency values are slightly higher compared to the movement of the same TdT::IFT81 measured in cultured procyclic parasites (Bertiaux et al., 2018b). This is possibly a consequence of the temperature increase likely to occur after mixing the cell suspension with warm liquid agar. Indeed, temperature can affect IFT rates and frequency as reported in trypanosomes (Buisson et al., 2013) and *Caenorhabditis elegans* (Jensen et al., 2018).

In conclusion, these results show that IFT speeds and frequencies are equivalent in cells with long and short flagella. Modulation of these parameters can therefore not explain the changes in flagellum length observed during *T. brucei* development.

### Unique IFT protein distribution in short epimastigote flagella

Immunofluorescence and live cell studies showed that IFT proteins were found in discrete spots along the flagellum and concentrated at the base of the flagellum in all trypanosome cell types. However, methanol fixation showed a lower abundance of the IFT pool at the flagellum base of short epimastigotes compared to the other stages (Fig. 4A & Fig. S1A). To understand whether it was real or if it was the mere consequence of fixation, live cells expressing the TdT::IFT81 were examined more closely. By eye, the IFT signal at the base indeed looked less intense but surprisingly, a rather large IFT pool was detected at the distal end of these short flagella (Fig. 6A & Video S4). This was investigated by quantitative analysis of the TdT::IFT81 fluorescent signal at the base; along the flagellum and at the distal tip in short epimastigote cells (n=17) using midgut trypomastigote cells as controls (n= 22). While the tip of the flagellum in control cells contained a reduced amount of IFT proteins (Fig. 6A-B), a really large IFT pool was detected at the distal tip of the short epimastigote flagellum (Fig. 6C) that turned out to be twice as abundant as the pool found at the base (Fig. 6D). This is in contrast to control midgut cells were the IFT signal at the tip was 4 to 5-fold lower compared to the base (Fig. 6B). This large amount of material could be engaged in IFT or simply be inactive, for example serving as a rapidly mobilizable pool to facilitate flagellum elongation in the salivary glands when these cells differentiate to produce the attached epimastigote stage with a longer flagellum (∼14 µm) (Rotureau et al., 2011).

**Fig. 6.**
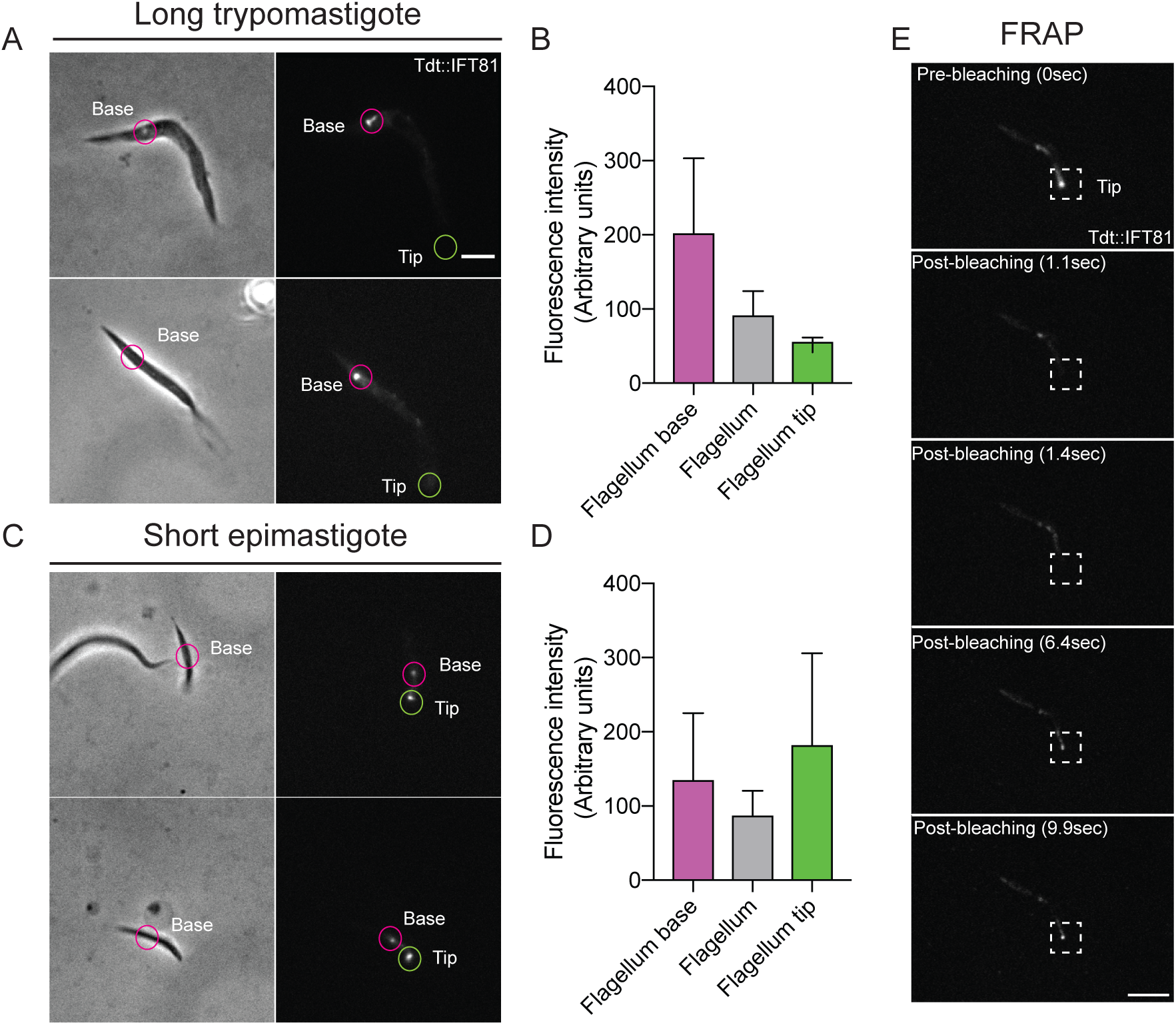
An IFT81 pool is present at the distal tip of short epimastigote flagella. Images of cells expressing a TdTomato-tagged version of IFT81 isolated from the tsetse fly cardia at the long trypomastigote stage (A) or at the short epimastigote stage (C). The phase contrast is shown on the right and the fluorescent signal is on the left. The base of the flagellum is indicated with a magenta circle and the tip with a green circle. The existence of an IFT pool at the tip is only detected in short epimastigote cells (also see Video S4 for live cells). Scale bar: 5 µm. (B, D) Quantification of the TdT::IFT81 mean fluorescence intensities at the base (magenta), in the middle region (grey) and the tip (green) of flagella in long trypomastigote (B, n=22) or in short epimastigote (D) cells (n=17). (E) FRAP analysis in a short epimastigote trypanosome expressing the TdT::IFT81 fusion protein. The distal end of the flagellum was bleached with a brief laser pulse and the fluorescence recovery was monitored during 20 sec. Pre-bleach situation: the IFT pool is present at the distal tip. Post-bleach situation: the signal becomes negative but rapid recovery was detected at the distal tip. The fluorescent signal is shown for the indicated times. The dotted square indicates the region around the distal tip. Scale bar: 3µm.

Fluorescence recovery after photobleaching (FRAP) was used to evaluate if this pool was dynamic. Short epimastigote cells were recovered from infected tsetse flies and a region of interest was drawn around the IFT signal at the distal tip for bleaching with a brief laser pulse in order to extinguish the fluorescent signal (Fig. 6E, second panel). Fresh anterograde trains were then detected, emerging from the basal pool, and traveling towards the tip of the flagellum (two examples are presented in Video S5). Within seconds, these trains reached the tip and progressively replenished the distal pool (Fig. 6E, bottom panels). Retrograde trains are frequently seen starting from this distal pool, demonstrating that this material is actively participating to IFT (Video S5). Such fluorescence recovery was observed at the distal tip of all the short flagella (n=7) that could be analysed.

### The length of the flagellum of short epimastigote cells does not increase after inhibition of cell division

In trypanosomes, the new flagellum is assembled whilst maintaining the existing one (Sherwin and Gull, 1989). This mature flagellum does not undergo turnover of its major axonemal components such as dynein arms or PFR components (Vincensini et al., 2018). This led us to propose a grow-and-lock model where the mature flagellum would be locked, preventing further elongation (Bertiaux and Bastin, 2020; Bertiaux et al., 2018b). In this context, two parameters cooperate to regulate length: the rate of elongation and the timing of the locking event that is linked to that of cell division. Manipulating one of these parameters resulted in the formation of flagella of different lengths (Bertiaux et al., 2018b).

Since IFT rate and frequency are equivalent in short and long flagella, one could consider that the short epimastigote cells have a tiny flagellum because of premature cell division, hence taking place before the flagellum could fully elongate. This was observed in procyclic cells where the amount of IFT kinesin had been knockdown by RNA interference: the number of IFT trains was significantly reduced, leading to the formation of flagella that were too short at the time of cell division. However, inhibition of cell division resulted in the construction of a flagellum that ultimately reached the same length as the parental flagellum (Bertiaux et al., 2018b). To address whether a modification of the timing of cell division could explain the production of flagella with different length in the context of the natural cyclical development of trypanosomes, cell division was chemically inhibited in parasites isolated from infected tsetse cardia. Cells were maintained for 24 hours in the presence of 10mM teniposide, a drug that interferes with mitochondrial DNA segregation but neither with basal body duplication nor with flagellum elongation (Robinson and Gull, 1991). In these conditions, cells cannot divide anymore. This arrests cell division and the impact on flagellum length can be analysed by IFA using DAPI to stain DNA and the axonemal marker Mab25 to measure flagellum length (Fig. 7). First, long trypomastigote parasites were analysed. These cells assemble a tiny flagellum before nucleus migration and transformation in epimastigote cells (Lemos et al., 2020). Remarkably, the flagellum elongated from ∼1 µm to an average of 6.2 ± 2.8 µm (n= 24)(Fig. 7, bottom left panels), a value that is even higher than the average flagellum length in short epimastigote cells (Rotureau et al., 2011). Surprisingly, the length of the old flagellum was shorter (21.4 ± 4.5 µm) than that of untreated cells (∼28 µm), suggesting possible remodelling. We then turned our attention to dividing epimastigote cells treated with teniposide. The new flagellum retained a length of 3.2 ± 1.2 µm and the old one of 27.9 ± 3.54 µm (n= 17)(Fig. 7, right panels). These values are comparable to those of untreated cells observed in this and other experiments (Fig. 1) and from previous *in vivo* studies (Rotureau et al., 2011; Sharma et al., 2008; Van Den Abbeele et al., 1999). Therefore, the short flagellum failed to elongate further, suggesting that its length is restricted.

**Fig. 7.**
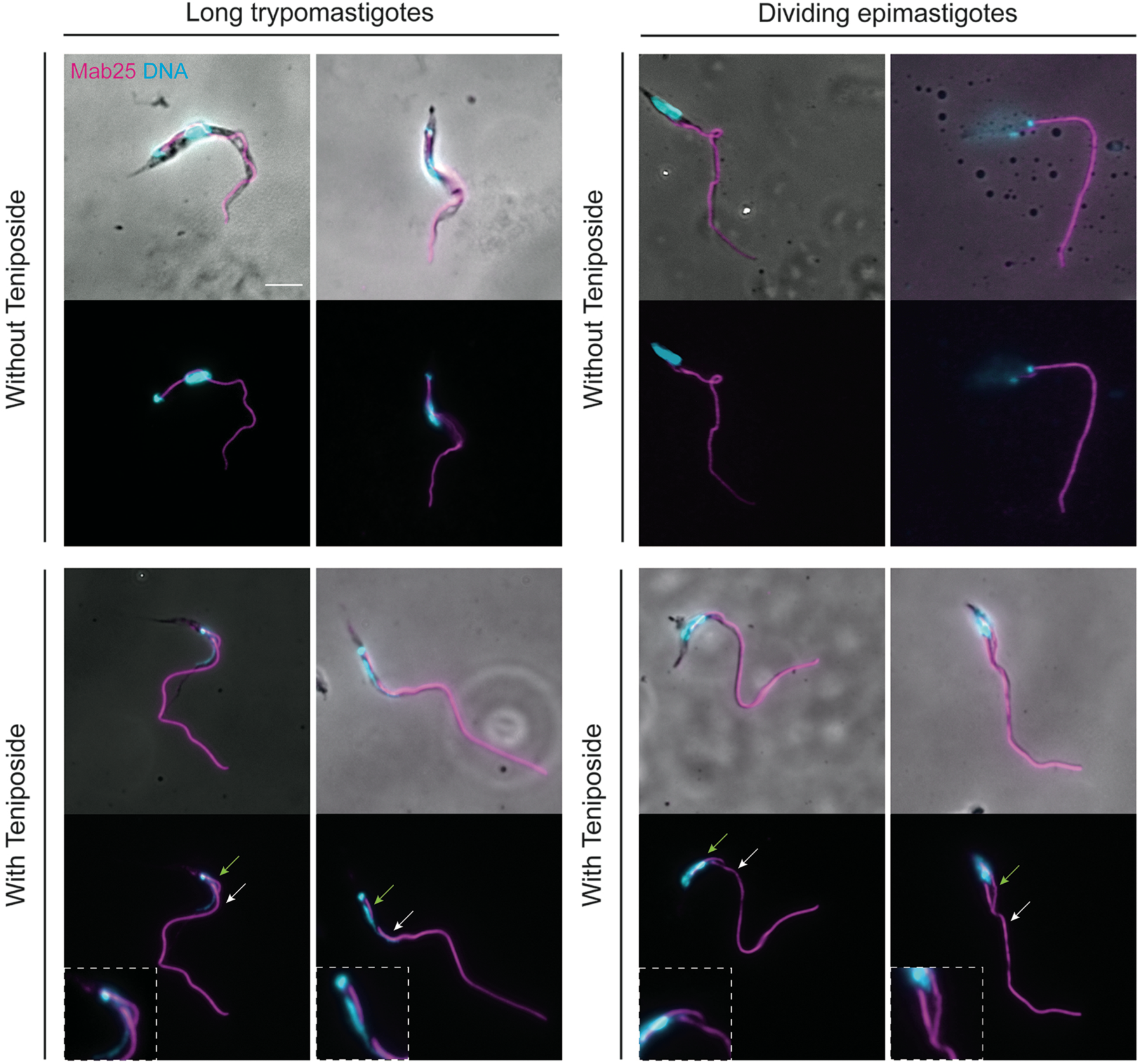
Inhibition of cell division has different impacts on flagellum length according to the trypanosome stage. Parasites isolated from tsetse cardia were treated (bottom panels) or not (top panels) with teniposide for 24 hours, fixed in methanol and stained with the Mab25 antibody to detect the axoneme (magenta) and DAPI to stain DNA (cyan). Left and right panels show untreated and treated cells, respectively. Two examples are shown for each situation. Green and white arrows show the new flagellum and the old flagellum, respectively. A significant increase of the length of the new flagellum is obvious in long trypomastigote cells whereas no visible impact could be detected for dividing epimastigote cells. Insets show higher magnification of the region with the new flagellum. Scale bar: 5 µm

## Discussion

IFT has been well studied in the procyclic stage of *T. brucei* in culture with a combination of functional and imaging studies. Here, transmission electron microscopy, FIB-SEM, immunofluorescence assays and live cell analysis revealed that IFT is present and active in all accessible parasite stages in the midgut, cardia and salivary glands of infected tsetse flies. The total amount of IFT proteins directly correlates with the length of each flagellum from 3 to 30 µm. A direct relationship between total amounts of IFT and final flagellum length was reported for the two equivalent flagella of *Chlamydomonas* (Marshall and Rosenbaum, 2001) and recently for the four pairs of flagella of different length in *Giardia lamblia* (McInally et al., 2019). There is however a significant difference between *Chlamydomonas* and *T. brucei*: in the first one the whole amount of IFT proteins is injected at once and train size progressively scales down leading to a reduction of the IFT concentration during elongation (Engel et al., 2009; Vannuccini et al., 2016) whereas in the second one, IFT recruitment to the flagellum progressively increases with length to maintain a constant concentration per unit of length (Bertiaux et al., 2018b).

Is there a causal relationship between amount of IFT and flagellum length? Reduction of IFT train frequency following knockdown of IFT kinesin using RNA interference led to the formation of shorter flagella (Bertiaux et al., 2018b), supporting the causal relationship. In *Chlamydomonas*, shifting the temperature-sensitive *fla10* kinesin mutant to intermediate temperature conditions resulted in a reduction in the number of IFT trains and in the formation of shorter flagella (Marshall and Rosenbaum, 2001). A simple model would be to postulate that flagellum length is controlled by the production of adequate amounts of IFT proteins. In this context, long and short epimastigote cells would synthesise more or less IFT proteins, respectively. However, this interpretation is difficult and should be taken with a pinch of salt because IFT is the construction machinery. Inhibition of cell division in kinesin RNAi conditions allowed trypanosomes to grow flagella twice as long without an increase in IFT trafficking (Bertiaux et al., 2018b). This model is further complicated by the existence of a large soluble pool of IFT proteins in the cytoplasm, reaching up to 50-fold the amount of IFT in the flagellum, at least in the case of *Chlamydomonas* (Ahmed et al., 2008; Wang et al., 2009). Because of the inherent difficulty to purify intact flagella in wild-type trypanosomes, this could not been quantified directly but IFA and observation of live cells expressing fluorescent IFT proteins indicate the presence of a substantial amount of IFT proteins in the cell body (Absalon et al., 2008).

Treatment of asymmetrically dividing epimastigote trypanosomes with teniposide showed that the short flagellum did not elongate further, which is in contrast with what was observed in the same experiment for long trypomastigote parasites coming from flies (Fig. 7) or procyclic trypanosomes in culture (Bertiaux et al., 2018b). Several reasons could explain this result. First, this flagellum could be locked prematurely. Addition of a cap, modification of microtubules or presence of a barrier at the base of the flagellum are potential mechanisms for such a locking model (Bertiaux and Bastin, 2020). Second, this flagellum could be constructed on the principle of the balance between assembly and disassembly, but with a marked disassembly rate as proposed in *Giardia* where increasing amounts of the microtubule-depolymerising kinesin 13 are associated with shorter flagellum length (McInally et al., 2019). The genome of *T. brucei* contains 5 genes for kinesin 13 and KIN13-2 is the only one to be localised at the distal tip of both mature and growing flagella (Chan et al., 2010). KIN13-2 could therefore be a candidate to limit elongation at the short epimastigote stage. However, functional investigation in procyclic cells in culture showed that the involvement of KIN13-2 on the control of flagellum length is marginal (Chan and Ersfeld, 2010). Third, the pool of cytoplasmic tubulin could be restricted as proposed in the limited soluble pool model (Goehring and Hyman, 2012). Intriguingly, the volume of the cytoplasm shrinks significantly at that stage (Rotureau et al., 2011) and a general reduction in cell activity has been observed (Natesan et al., 2007). In these conditions, IFT could remain active but if there is no or little tubulin transport, elongation could not continue further (Craft et al., 2015).

In the fly, it is commonly accepted that the short epimastigote cell is the one that establishes the infection in the salivary glands and that transforms in the attached epimastigote stage (Van Den Abbeele et al., 1999). This stage is characterised by a longer flagellum (14 µm) that forms multiple membrane protrusions for adhesion to the epithelium of the salivary glands (Tetley and Vickerman, 1985). This means that the mechanisms restricting flagellum length must be abolished to allow further elongation. Based on the models proposed above, the flagellum could be unlocked, the activity of a depolymerising kinesin could be inhibited or the synthesis of tubulin and other flagellar precursors could resume. Finally, given the difficulty to visualise the establishment of the infection by a single parasite in the salivary glands of the tsetse fly, construction of a new, longer, flagellum followed by an asymmetric division cannot be formally excluded. Such a situation has been observed for the differentiation of the attached epimastigote to the trypomastigote stage preceding the transformation in metacyclic parasites that are released in the saliva (Rotureau et al., 2012).

What processes could govern the formation of a longer flagellum? This is observed during the differentiation from the trypomastigote procyclic stage to the epimastigote stage in the cardia (Rotureau et al., 2011; Sharma et al., 2008; Van Den Abbeele et al., 1999). We propose that the new flagellum inherited from the last division of the procyclic stage is not locked and therefore can therefore elongate further giving rise to the long trypomastigote cells encountered in the cardia. If this flagellum is not locked, it must be more dynamic, meaning it could explain its unexpected shortening following inhibition of cell division while the new flagellum elongates from 1 to 6 µm (Fig. 7). This suggests that the total amount of tubulin might be limiting, giving support to the soluble pool model as a possible mode of control of flagellum length. Further experimentation could address these models but would be greatly helped by an *in vitro* cultivation system. The overexpression of the RNA binding protein 6 (RBP6) recapitulates some aspects of trypanosome development in the tsetse fly including the production of metacyclic cells that are responsible for infection of mammalian hosts (Kolev et al., 2012). Unfortunately, it bypasses most intermediate stages, including the asymmetrically dividing epimastigote stage, and cells with short or long flagella have not been reported (Kolev et al., 2012).

Like multicellular organisms, trypanosomes have the ability to assemble flagella of different length. This is likely to be achieved by different mechanisms: if the grow-and-lock model can explain the situation encountered in procyclic trypanosomes, it does not seem to apply to the situation of dividing epimastigote cells, where a different process could be involved. In the related parasites *Trypanosoma congolense* and *Trypanosoma vivax*, shortening of existing flagella was reported (Peacock et al., 2018) or suspected (Ooi et al., 2016). This feature has so far not been reported for the life cycle of *T. brucei*. However, it could be triggered here in experimental conditions for long trypomastigote cells present in the cardia, suggesting that there might be more hidden features in the control of flagellum length in trypanosomes. Overall, these results highlight the importance of studying different flagellated organisms as it can bring findings reflecting the diversity of cilia across evolution but also within single organisms such as humans.

## Acknowledgements

We thank Aline Crouzols and Christelle Travaillé for help with tsetse fly infection and dissection; Moara Lemos for help with sample preparation and Derrick Robinson (University of Bordeaux) for providing the Mab25 antibody. We are grateful to the Photonic Bioimaging and Ultrastructural Bioimaging facilities for access to their equipment, and to Gérard Pehaud-Arnaudet for training in electron microscopy.

## Author contributions

Conceptualization: E.B, B.R., P.B.; Methodology: E.B., A.M.; Formal analysis: E.B., A.M..; Writing – original draft: E.B., P.B.; Writing - review & editing: E.B., A.M., B.R., P.B.; Supervision: B.R., P.B.; Funding acquisition: E.B., P.B.

## Legend of supplementary figures

**Fig. S1.**
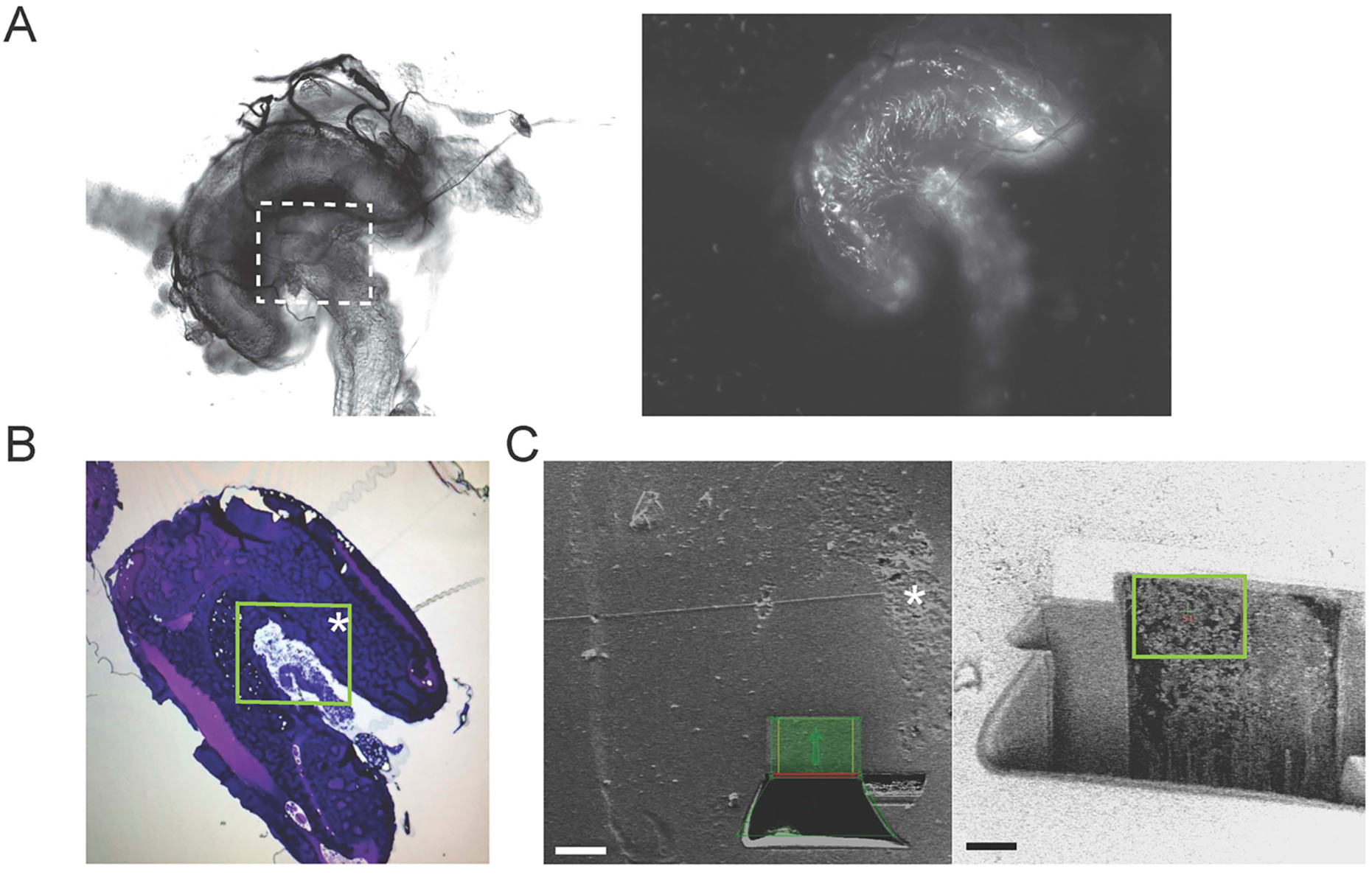
Use of a correlative light and electron microscopy approach to detect trypanosomes in infected tsetse cardia. (A) Entire cardia isolated from a tsetse fly infected with trypanosomes expressing a TandemTomato fluorescent reporter (Calvo-Alvarez et al., 2018). Left panel: Brightfield image showing the general morphology of the cardia. The square indicates the region of interest for electron microscopy. Right panel: The same cardia viewed by fluorescence microscopy reveals the presence of trypanosomes in various positions within the organ. (B) A semi-thin section of the cardia stained with toluidine blue. Parasites are visible in the green square. The same region of interest was observed on the semi-thin section and on the block surface by scanning electron microscopy (white stars on B and C). (C) Different views of the bloc surface of the sample analysed by FIB-SEM after a cross-sectioning and a polishing by FIB. Parasites are visible on the side view (green squares). Scale bars: 20 µm (left) and 10 µm (right).

**Fig. S2.**
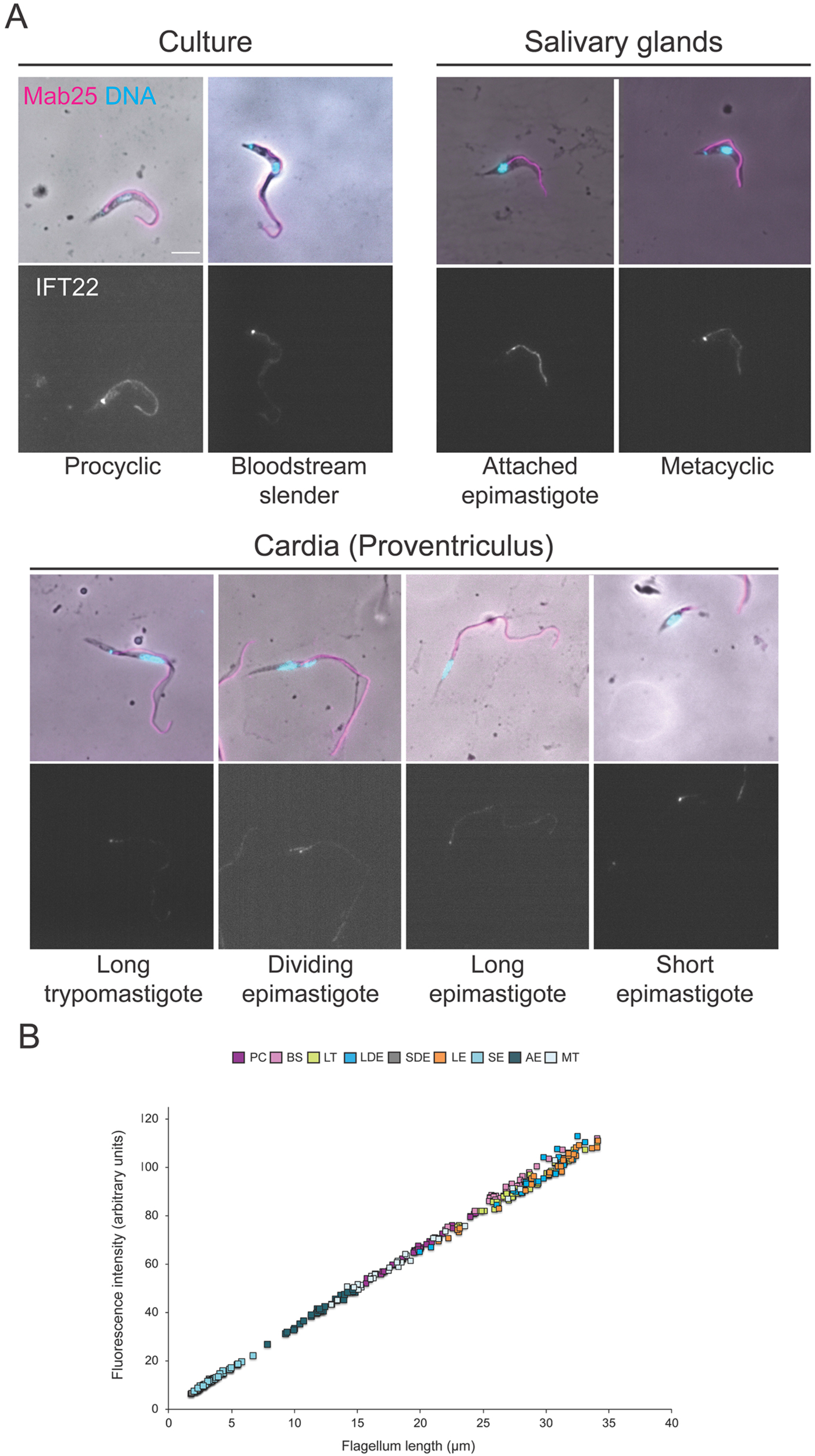
IFT22 distribution during the *T. brucei* parasite cycle. (A) Parasites isolated from tsetse flies or grown in culture were fixed in cold methanol and stained with the Mab25 antibody to detect the axoneme (magenta) and the anti-IFT22 antibody (white). The top panels show the phase-contrast images merged with DAPI (cyan) and Mab25 signals (magenta). The bottom ones show the IFT22 fluorescent signal (white). Scale bar: 5µm. (B) Quantification of the total amount of IFT22 fluorescent signal in the flagellum normalized to flagellum length for each stage of the parasite cycle. The ROI was defined by the axonemal marker and used to measure the flagellum length. The fluorescence intensity directly reflects the total amount of IFT22 proteins present in the flagellum compartment. n=35 cells per stage. BSF: Slender bloodstream form, PC: Procyclic form, LT: Long trypomastigote, LDE: Long dividing epimastigote, SDE: Short dividing epimastigote, LE: Long epimastigote, SE: Short epimastigote, AE: Attached epimastigote, MT: Metacyclic trypomastigote.

## Supplementary videos

**Video S1**. FIB-SEM analysis of parasites in the cardia. The region shown corresponds to that indicated by black squares at Figure 2. Two IFT particles are indicated by arrowhead: the first one (IFT1) is a long train found on doublet 8 and the second one is present on doublet 4 (IFT2). This video is related to Figure 2.

**Video S2**. IFT trafficking in bloodstream (first in sequence) and procyclic cells grown in culture. Cells are expressing TdT::IFT81 following *in situ* tagging and display IFT trafficking. The bright signal at the base of the flagellum is clearly visible. The video plays at real time. This video is related to Figure 5.

**Video S3**. IFT trafficking in parasites isolated from the salivary glands showing first an attached epimastigote and then a metacyclic cell. Trypanosomes are expressing TdT::IFT81 following *in situ* tagging and display IFT trafficking. The bright signal at the base of the flagellum is clearly visible. The video plays at real time. This video is related to Figure 5.

**Video S4**. IFT trafficking in parasites isolated from the cardia of an infected tsetse fly. The sequence shows successively a long trypomastigote, a dividing epimastigote (trafficking visible in both old and new flagella), a long epimastigote (retrograde trains are nicely visible at the end of this sequence) and a short epimastigote with the typical accumulation of IFT proteins at the tip of the short flagellum. Cells are expressing TdT::IFT81 following *in situ* tagging and display IFT trafficking. The bright signal at the base of the flagellum is clearly visible. The video plays at real time. This video is related to Figure 5.

**Video S5**. Two examples of FRAP analysis of short epimastigote cells expressing TdT::IFT81. The distal portion of the flagellum was bleached with a brief laser pulse. Entry of new anterograde IFT trains in the flagellum can be monitored, they reach the distal tip where signal progressively comes backs before smaller, less bright, retrograde trains can be detected. This video is related to Figure 6. The first cell goes briefly out of the plane of focus.

